# GM_1_-oligosaccharide rescues rotenone-impaired neuronal polarization through RhoA/ROCK modulation and mitochondrial protection

**DOI:** 10.64898/2025.12.29.696900

**Authors:** Pablo E. A. Rodríguez, Guillermo N. Colmano, Andrea Pellegrini, María E. Mariani, Silvana B. Rosso, Gonzalo Quassollo, Pablo Helguera, Mariano Bisbal, Gerardo D. Fidelio, Mónica S. Sanchez

## Abstract

Neuronal polarization is a fundamental process in the formation of functional neural circuits, relying on the precise coordination between cytoskeletal regulatory signals and mechanisms that sustain cellular integrity. Disruption of these processes compromises neuronal differentiation and survival, and various neurotoxic compounds, including certain pesticides, have been associated with such dysfunctions. In this context, identifying molecules that counteract these detrimental effects is of significant therapeutic interest.

Neuronal polarization is essential for the establishment of functional neural circuits and relies on coordinated regulation of actin cytoskeleton dynamics, RhoA/ROCK signaling, and mitochondrial function. Here, we investigated the neuroprotective and neurorestorative potential of the ganglioside GM1 and its oligosaccharide derivative, osGM1, in primary hippocampal pyramidal neurons exposed to the mitochondrial neurotoxin rotenone.

Rotenone induced a marked arrest of neuronal development, impaired axonal elongation, and disrupted mitochondrial organization and membrane potential. Both GM1 and osGM1 promoted recovery of neuronal polarity and axonal growth, exerting protective and restorative effects even under continuous toxin exposure, with osGM1 showing superior efficacy. Notably, osGM1 also reversed axonal growth deficits caused by pathological actin stabilization. Mechanistically, osGM1 normalized rotenone-induced hyperactivation of the RhoA/ROCK pathway without altering basal signaling and partially restored mitochondrial network integrity and function.

Collectively, these findings identify osGM1 as a multi-target modulator of cytoskeletal and mitochondrial dysfunction and support its translational potential as a therapeutic strategy to counteract neurotoxin-induced neuronal damage.

## Introduction

Neuronal polarization is a fundamental process required for the establishment of functional neural circuits. This process depends on the coordinated regulation of multiple cellular events, particularly the RhoA/ROCK pathway, actin cytoskeleton dynamics, and mitochondrial integrity (Banker, 2018; Smith & Gallo, 2018; Takano, Funahashi, & Kaibuchi, 2019). Because these mechanisms are tightly interconnected, an initial disruption in polarity regulators can trigger a cascade of alterations that ultimately compromise neuronal viability.

Among the agents capable of disturbing this sequence, various neurotoxic compounds—including pesticides and mitochondrial toxins such as rotenone—have been shown to interfere with neuronal polarization (Bisbal, Remedi, Quassollo, Caceres, & Sanchez, 2018; Bisbal & Sanchez, 2019; Fazzari et al., 2023; Sanchez, Gastaldi, Remedi, Caceres, & Landa, 2008). Notably, rotenone activates the RhoA/ROCK pathway, thereby altering the early mechanisms that govern neuronal polarity (Bisbal et al., 2018). This aberrant activation of RhoA/ROCK directly impacts actin cytoskeleton dynamics, a key determinant of axon specification and neurite elongation (Da Silva et al., 2003; Dupraz et al., 2019; Wojnacki et al., 2024). Moreover, because actin organization modulates axonal transport and subcellular organelle distribution (Chada & Hollenbeck, 2004; Hollenbeck & Saxton, 2005; Mandal & Drerup, 2019; Smith & Gallo, 2018), actin defects can secondarily—but critically—affect mitochondrial function, amplifying the initial damage caused by the toxin.

The relevance of this polarity-RhoA/ROCK-actin-mitochondria axis becomes even clearer considering that mitochondria are also a primary target of rotenone, which inhibits complex I and induces energetic dysfunction (Greenamyre, Betarbet, & Sherer, 2003). Thus, this toxin exerts a dual impact: direct bioenergetic impairment plus secondary cytoskeleton-dependent mitochondrial disturbances.

In this scenario, identifying molecules that can interrupt this pathological sequence and preserve neuronal integrity is crucial. Gangliosides, a class of glycosphingolipids abundant in the nervous system, have attracted considerable interest because of their role in regulating essential processes for neuronal function and survival (Svennerholm, 1980; Ledeen & Wu, 2018). Among them, the ganglioside GM1 stands out for its roles in neuronal differentiation, signaling, synaptic plasticity, and neuroprotection (Aureli et al., 2016; Chiricozzi et al., 2020; Lunghi et al., 2020; Schengrund & Prouty, 1988; Schengrund & Shochat, 1988; Wieraszko & Seifert, 1985). GM1 is mainly located in the outer leaflet of the plasma membrane, where it modulates essential signaling cascades and interacts with neurotrophic factor receptors, including TrkA, TrkB, and TrkC, coordinating molecular events that determine neuronal fate (Hansson, Holmgren, & Svennerholm, 1977; Mocchetti, 2005; Rabin & Mocchetti, 1995).

Recent studies have shown that the oligosaccharide chain of GM1 (osGM1) is essential for its neurotrophic properties. This fragment activates pathways that promote neuronal differentiation and protects neurons from harmful insults (Chiricozzi et al., 2019; Chiricozzi et al., 2021; Chiricozzi et al., 2017). In addition, GM1 and osGM1 counteract the effects of various neurotoxic substances and oxidative stress, reinforcing their role as guardians of neuronal integrity (Aureli et al., 2016; Vlasova Iu, Zakharova, Sokolova, & Avrova, 2013; Wang, Li, Yu, Zhou, & Gao, 2022). In this context, understanding whether these gangliosides can modulate the RhoA/ROCK pathway and stabilize the actin cytoskeleton is key to determining whether they can block the degenerative cascade that ultimately compromises mitochondrial function.

Previous studies from our group demonstrated that rotenone alters RhoA/ROCK signaling, contributing to axonal impairment (Bisbal et al., 2018; Bisbal & Sanchez, 2019), while independent research has shown that rotenone directly impairs mitochondrial activity (Greenamyre et al., 2003). Together, these findings highlight the importance of evaluating whether GM1 and osGM1 can counteract this toxic sequence by maintaining neuronal polarity, preserving actin organization, and protecting mitochondrial function.

In the present study, we investigated the neuroprotective and neurorestorative effects of GM1 in primary cultures of hippocampal pyramidal neurons exposed to rotenone. We aimed to determine the ability of both compounds to prevent or reverse rotenone-induced alterations in neuronal polarity, cytoskeletal organization, and mitochondrial activity. Our results show that GM1 and osGM1 support neuronal polarity and axonal growth under toxic conditions, and that osGM1 displays superior efficacy in restoring neuronal morphology and preserving mitochondrial function, underscoring its potential as a neuroprotective and neurorestorative molecule.

## Methods

### Animals use and care

Pregnant Wistar rats were born in the vivarium of INIMEC-CONICET-UNC (Córdoba, Argentina). Wistar rat lines were originally provided by Charles River Laboratories International Inc (Wilmington, USA). All procedures and experiments involving animals were approved by the Animal Care and Ethics Committee (CICUAL http://www.institutoferreyra.org/en/cicual-2/) of INIMEC-CONICET-UNC (Resolution numbers 014/2017 B, 015/2017 B, 006/2017 A and 012/2017 A) and were in compliance with approved protocols of the National Institute of Health Guide for the Care and Use of Laboratory Animals (SENASA, Argentina).

All methods were carried out in accordance with relevant guidelines and regulations.

### GM1 Ganglioside and GM1 Oligosaccharide (osGM1)

GM1 ganglioside from porcine brain and GM1 oligosaccharide, prepared by ozonolysis followed by alkaline degradation of GM1 as described by Wiegandt & Bücking (1970), were generously gifted by TRB Pharma S.A. (Buenos Aires, Argentina). Altogether, nuclear magnetic resonance, mass spectrometry, and HPTLC analyses showed greater than 99% homogeneity for both GM1 ganglioside and GM1 oligosaccharide.

### Neuronal Cultures

Primary hippocampal cultures were prepared as previously described (Bisbal et al., 2018). Briefly, hippocampi were dissected from E18 rat embryos, incubated in 0.25% trypsin (Thermo Fisher Gibco; Cat. No. 15090-046) for 15 min at 37 °C and mechanically dissociated using a Pasteur pipette. Cells were plated on 12-mm glass coverslips (Marienfeld Superior; Cat. No. 633029) pre-coated with poly-L-lysine (1 mg/mL; Sigma; Cat. No. P2636) at a density of 2000 cells/cm².

Cultures were maintained in MEM (Thermo Fisher Gibco; Cat. No. 61100-061) supplemented with penicillin–streptomycin (Cat. No. 15140122), CTS GlutaMAX I (Cat. No. A1286001), sodium pyruvate (Cat. No. 11360070), and 10% horse serum (Cat. No. 16050122). After 2 h, coverslips were transferred to Neurobasal medium supplemented with B-27 Plus (Cat. No. A3582801) and GlutaMAX I.

### Drugs and Pharmacological Treatments

Rotenone (Sigma-Aldrich; Cat. No. R8875, PubChem CID: 6758) was used at a concentration of 0.1 µM, which has been shown not to induce apoptosis (Sanchez et al., 2008). Jasplakinolide (1 nM; Sigma-Aldrich; Cat. No. J4580, PubChem CID: 9831636) was used as described by Alvariño et al. 2021 (Alvarino et al., 2021). Both compounds were dissolved in dimethyl sulfoxide (DMSO, 0.01%; Sigma-Aldrich, Cat. No. D5879), and control cultures received an equivalent amount of DMSO. GM1 and osGM1 were used at 70 µM. Neurons were treated with DMSO for 48 h, rotenone for 48 h, GM1 for 48 h, osGM1 for 48 h, or with sequential combinations (rotenone for 24 h → osGM1 for 24 h, or osGM1 for 24 h → rotenone for 24 h). In all combined treatments, the medium was not replaced between treatments, allowing both compounds to remain present during the final 24 h corresponding to the second treatment.

### Immunofluorescence

Cells were fixed at room temperature with 4% paraformaldehyde (Sigma-Aldrich; Cat. No. 441244) in PBS containing 4% sucrose for 20 min. After PBS washes, samples were permeabilized with 0.2% Triton X-100 (Bio-Rad; Cat. No. 1610407) for 5 min and blocked for 1 h in 5% BSA. Primary antibody incubations were performed for 1 h at 24 °C, followed by 1 h at 37 °C with Alexa 488 or Alexa 568 secondary antibodies (Molecular Probes; 1:1000). Coverslips were mounted in FluorSave (Millipore Calbiochem). F-actin was labeled with rhodamine-phalloidin (1:1500). A mAb against tyrosinated-α-Tubulin (Tyr-Tub; Millipore Sigma Antibody; Cat Number: MAB1864, RRID: AB_2210391) was used at 1:1000.

### Morphometric Analysis of Neuronal Parameters

Morphometric analyses were performed on maximum-intensity projections acquired on a Zeiss LSM 800 confocal microscope. Neurite length was quantified in antibody-labeled neurons randomly selected and manually traced following established procedures (Bisbal et al., 2018; Sanchez et al., 2008). Images were processed in Fiji/ImageJ (Schindelin et al., 2012).

### DNA constructs

The following cDNA constructs were used in this study: cDNA coding for unimolecular biosensors for RhoA (RhoA2G; (Fritz et al., 2013) and ROCK (Eevee-ROCK; (C. Li et al., 2017) were kindly provided by Drs. Olivier Pertz and Michiyuki Matsuda to Dr. Bisbal. The RhoA2G biosensor uses mTFP/mVenus as acceptor/donor fluorophores pair, whereas ROCK uses CFP/YFP; in both cases, FRET increases upon RhoA-GTP activation or ROCK-mediated phosphorylation. A cDNA coding for the mitochondrial-targeted yellow fluorescent protein (pCAG-mitoYFP; (Zamponi et al., 2018).

### Transient Electroporation

Primary hippocampal neurons were electroporated using the Nucleofector II system (Lonza; Cat. No. AAD-1001N) according to the manufacturer’s protocol (program O-003, optimized for primary hippocampal neurons). 0.5×10^6^ - 1×10^6^ cells were used for each transfection with 3 µg of plasmid DNA (pCAG-mitoYFP, RhoA biosensor or Eevee-ROCK).

### FRET Imaging of RhoA and ROCK Activity

RhoA and ROCK activity were assessed using FRET ratiometric analysis as previously described (Bisbal et al., 2018; Wojnacki et al., 2024). Images were acquired on a Zeiss LSM 980 equipped with Airyscan 2. The donor (cyan/teal) was excited at 458 nm, while donor and acceptor emissions were collected simultaneously. For ratio imaging FRET calculation, the donor channel (donor emission) and FRET channel (acceptor emission) images were smoothed with a median filter (1.5 pixel ratio), background subtracted (50.0 pixels rolling ball radius) and aligned. FRET map images were generated by dividing the processed FRET channel images over donor channel images. To remove out-of-cell pixels from the analysis, a 0–1 intensity value binary mask was created using the FRET channel images and multiplied by the FRET map images. Finally, pixel values of the FRET maps images were color coded using a custom look-up table (LUT). All image processing was coded in an ImageJ macro so all images were processed in the same way (available upon request).

### Mitochondrial Membrane Potential

Mitochondrial membrane potential was measured with MitoTracker Red CM-H₂XRos (Thermo Fisher Scientific), following Hao et al. 2022 (Hao et al., 2022). After 48 h of treatment, as described in Section 2.3, neurons were incubated with the 100 nM dye for 20 min at 37 °C. Images were obtained on a Zeiss LSM 800, and fluorescence intensity was quantified in Fiji/ImageJ as a readout of mitochondrial potential.

### Mitochondrial Morphometric Analysis

Pyramidal neurons expressing mitoYFP were used to visualize mitochondria by fluorescence microscopy. After 48 h, cells were fixed and images analyzed in Fiji/ImageJ. Fluorescence images were converted to grayscale and binarized to segment individual organelles. The Analyze Particles tool was used to extract morphometric parameters: maximum (Feret) and minimum (MinFeret) diameters, area, and perimeter. The Feret diameter, defined as the maximum distance between parallel tangents of an object’s contour, was used as a robust measure of mitochondrial length (Buchanan et al., 2023; Fischer, Dash, Liu, & Waxham, 2018).

Mitochondria were categorized as: punctate (Feret ≤ 300 nm), globular (Feret 600–1200 nm), or tubular (Feret ≥ 500 nm and MinFeret < 0.7 × Feret)(Helguera et al., 2013). Organelles not fitting these ranges were grouped as morphologically indeterminate.

### Statistical Analyses

Data are presented as mean ± SEM. Statistical comparisons were performed using one-way ANOVA followed by Duncan’s test (p < 0.05). Normality was verified using the Shapiro-Wilk test. Analyses were conducted in InfoStat using generalized linear models (Di Rienzo et al., 2012).

## Results

### osGM1 enhances neuronal polarization and reverses the rotenone-induced deficit

Hippocampal pyramidal neurons in culture follow a highly ordered morphological program before developing a single axon and several short dendrites. After plating, cells exhibit no evident polarity and are characterized by broad lamellipodia (Stage 1). They subsequently extend several similar processes (minor neurites) at Stage 2. Finally, one of these differentiates into the axon through a prominent, dynamic growth cone enriched in labile actin and unstable microtubules (Stage 3). Days later, the remaining processes differentiate into dendrites (Banker, 2018). This predictable sequence provides a robust framework for evaluating how different treatments modulate progression toward neuronal polarization.

In this context, the effects of GM1, osGM1, rotenone, and their combinations on neuronal polarity were analyzed using a one-way ANOVA, which revealed highly significant differences among groups (F(7,40) ≈ 2704.93, p ≤ 0.0001). As previously described, rotenone induces a developmental arrest of hippocampal pyramidal neurons at Stage 2 ((Sanchez et al., 2008); Fig. 1B, I). While most control cells reached Stage 3 (Fig. 1A, B), neurons exposed to the toxin remained stalled at Stage 2. In contrast, both GM1 (Fig. 1C, I) and osGM1 (Fig. 1D, I) markedly increased the proportion of neurons achieving polarization, with osGM1 emerging as the most effective treatment.

**Figure 1.**
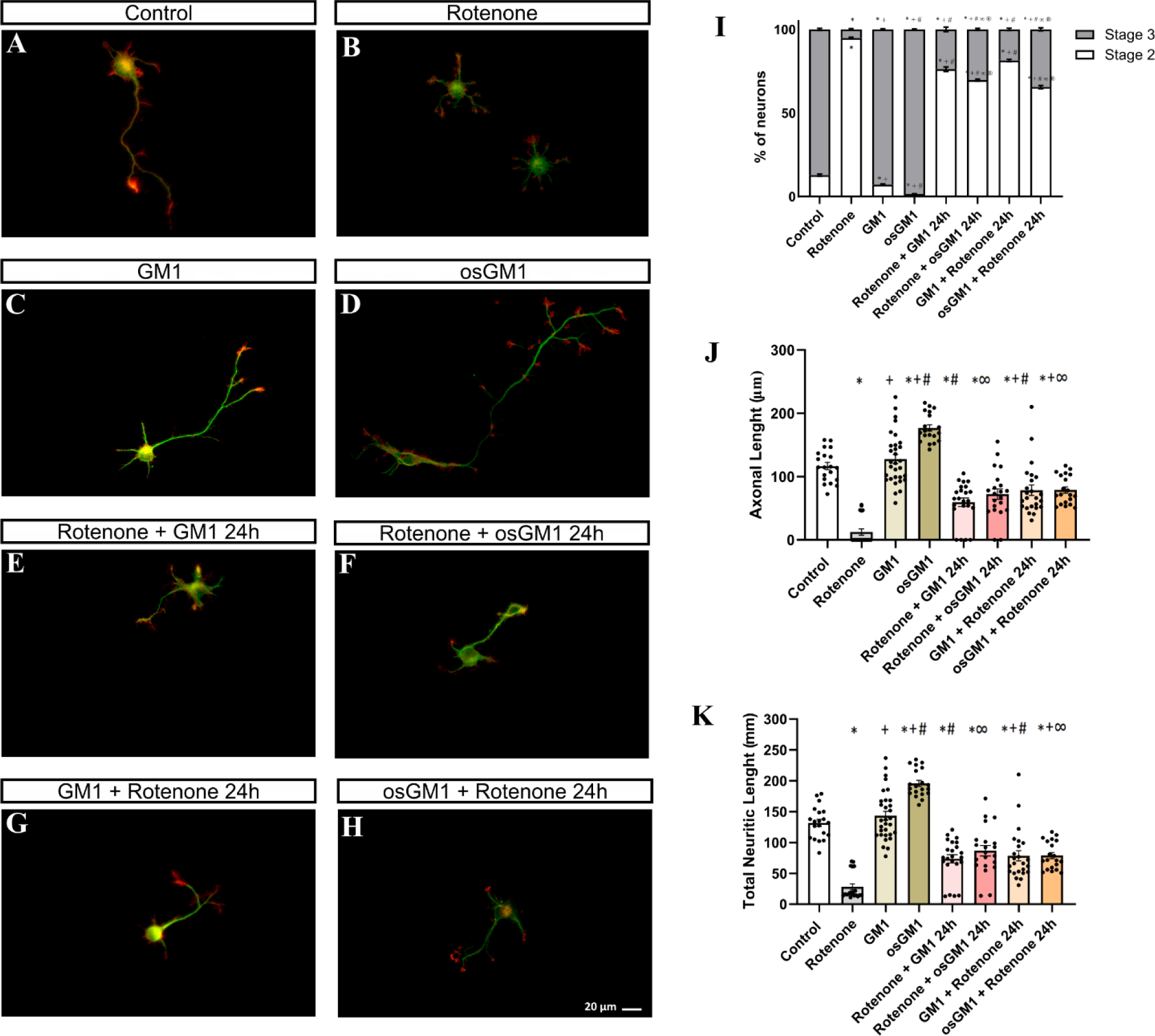
GM1 and osGM1 promote neuronal polarization and rescue rotenone-induced deficits in hippocampal neurons. (A-H) Representative images of hippocampal neurons under control conditions or treated with rotenone, GM1, osGM1, or their combinations using neuroprotective or neurorestorative paradigms. Neurons were stained with mAb against tyrosinated α-tubulin (green) and phalloidin-Rhodamine (red). Scale bar: 20 μm. (I) Graph showing the percentage of stage 2 and stage 3 neurons. The percentages were calculated from six independent cultures. (J, K) Graphs showing quantification of axonal length (J) and total neurite length (K) per neuron. Rotenone induced a developmental arrest at Stage 2 and impaired axonal elongation and neuritic growth. GM1 and osGM1 promoted neuronal polarization and reversed these deficits under both paradigms, with osGM1 showing greater efficacy. For all quantitative analyses, 20–30 neurons were analyzed per treatment condition. Graphs represent mean ± SEM. Statistical analysis: one-way ANOVA followed by Duncan’s post hoc test (p < 0.05). Significant differences versus control (*), rotenone-treated (+), GM1-treated (#), and the corresponding GM1-treated condition (∞).

To evaluate the neurorestorative potential of the gangliosides, it was considered that neurons can partially recover their polarization upon rotenone removal (Sanchez et al., 2008). In our study, however, rotenone remained present throughout the 48 h treatment period, and gangliosides were added only after the first 24 h. Under these conditions of continuous toxin exposure, the observed recovery reflects the ability of GM1 (Fig. 1E, I) or osGM1 (Fig. 1F, I) to reverse rotenone-induced toxicity. Likewise, when GM1 (Fig. 1G, I) or osGM1 (Fig. 1H, I) were present from the start of the experiment, and rotenone was added only during the final 24 h, both compounds exerted an apparent neuroprotective effect, preventing disruption of the normal progression toward neuronal polarization. Under both conditions -neurorestoration and neuroprotection- GM1 and, notably, osGM1 restored progression to neuronal polarization, with osGM1 displaying the highest efficacy.

The impact of the treatments on neuritic growth was assessed using a one-way ANOVA, which revealed significant differences in axonal length (F(9,160) = 45.99, p ≤ 0.0001) and total neurite length per neuron (F(9,160) = 49.98, p ≤ 0.0001). As previously demonstrated, rotenone markedly reduced axonal length (Fig. 1B, J) relative to the control (Fig. 1A, J) and significantly reduced total neurite length (Fig. 1B, K) relative to the control (Fig. 1A, K), consistent with reports describing truncated or absent axons in Stage 3 neurons (Sanchez et al., 2008). In contrast, the presence of GM1 throughout the 48 h treatment significantly increased axonal length (Fig. 1C, J) and total neurite length (Fig. 1C, K). Similarly, the presence of osGM1 throughout the 48 h treatment significantly increased axonal length (Fig. 1D, J) and total neurite length (Fig. 1D, K), with osGM1 producing the most robust effects.

Moreover, under the scheme described above -leaving the first compound in the medium for the entire 48 h and adding the second only during the final 24 h- two complementary experimental approaches were distinguished. The late addition of GM1 or osGM1 in the continuous presence of rotenone enabled neurorestoration to be assessed. Conversely, the late addition of the toxin to neurons previously exposed to gangliosides enabled neuroprotection to be evaluated. In both conditions, GM1 -whether added before or after rotenone- reduced the toxin-induced inhibition of axonal elongation (Fig. 1G, J and Fig. 1E, J, respectively) and total neurite growth (Fig. 1G, K and Fig. 1E, K, respectively), without altering the length of minor processes, which maintained their relative proportion to the axon. Similarly, osGM1 -whether added before or after rotenone- reduced the toxin-induced inhibition of axonal elongation (Fig. 1H, J and Fig. 1F, J, respectively) and total neurite growth (Fig. 1H, K and Fig. 1F, K, respectively), without altering the length of minor processes, which maintained their relative proportion to the axon.

Altogether, the data presented in figure 1 indicate that GM1 and osGM1 not only protect neurons from the toxic effect of rotenone on axonal elongation but also promote recovery once damage has been established. The superior ability of osGM1 to reverse these alterations is particularly noteworthy. Given that gangliosides were the most effective treatments for restoring polarization, the next step was to evaluate whether they also normalized actin cytoskeletal dynamics, a central component of the transition to Stage 3.

### osGM1 modulates actin cytoskeleton dynamics and promotes axonal elongation

Neuronal polarization critically depends on actin remodeling within the growth cone. Numerous studies in hippocampal neurons have demonstrated that axonal extension requires high actin dynamics, proper microtubule stabilization, and a finely orchestrated interplay among membrane lipids, proteins, and the extracellular matrix (Bradke & Dotti, 1999; Buck & Zheng, 2002; Pfenninger, 2009; Sonnino, Mauri, Chigorno, & Prinetti, 2007; Witte, Neukirchen, & Bradke, 2008). Since in the above experiments we identified that osGM1 act as the most effective compound for promoting polarization, subsequent experiments were designed to determine whether its action is associated with the ability to counteract aberrant actin stabilization, a central mechanism underlying rotenone-induced blockade (Sanchez et al., 2008). To this end, we evaluated whether osGM1 can reverse deficits specifically arising from reduced actin dynamics using Jasplakinolide, a persistent actin filament stabilizer (Holzinger, 2001).

To assess whether osGM1 can reverse the effects of prolonged actin stabilization, we first compared control neurons (Fig. 2A), neurons treated with jasplakinolide for 72 h (Fig. 2B), and neurons treated with jasplakinolide and subsequently co-exposed to osGM1 during the last 48 h (Fig. 2C). ANOVA revealed significant differences in both axonal length (F(2,91) = 23.64, p ≤ 0.0001) and total neuritic length (F(2,91) = 21.98, p ≤ 0.0001) (Fig. 2A-E). Jasplakinolide (1 nM) caused a significant reduction in axonal growth and total neuritic length compared with controls (p ≤ 0.05). In contrast, co-treatment with osGM1 during the last 48 h attenuated this inhibition. It significantly increased both axonal and total neuritic length compared with treatment with jasplakinolide alone, and even relative to control conditions.

**Figure 2.**
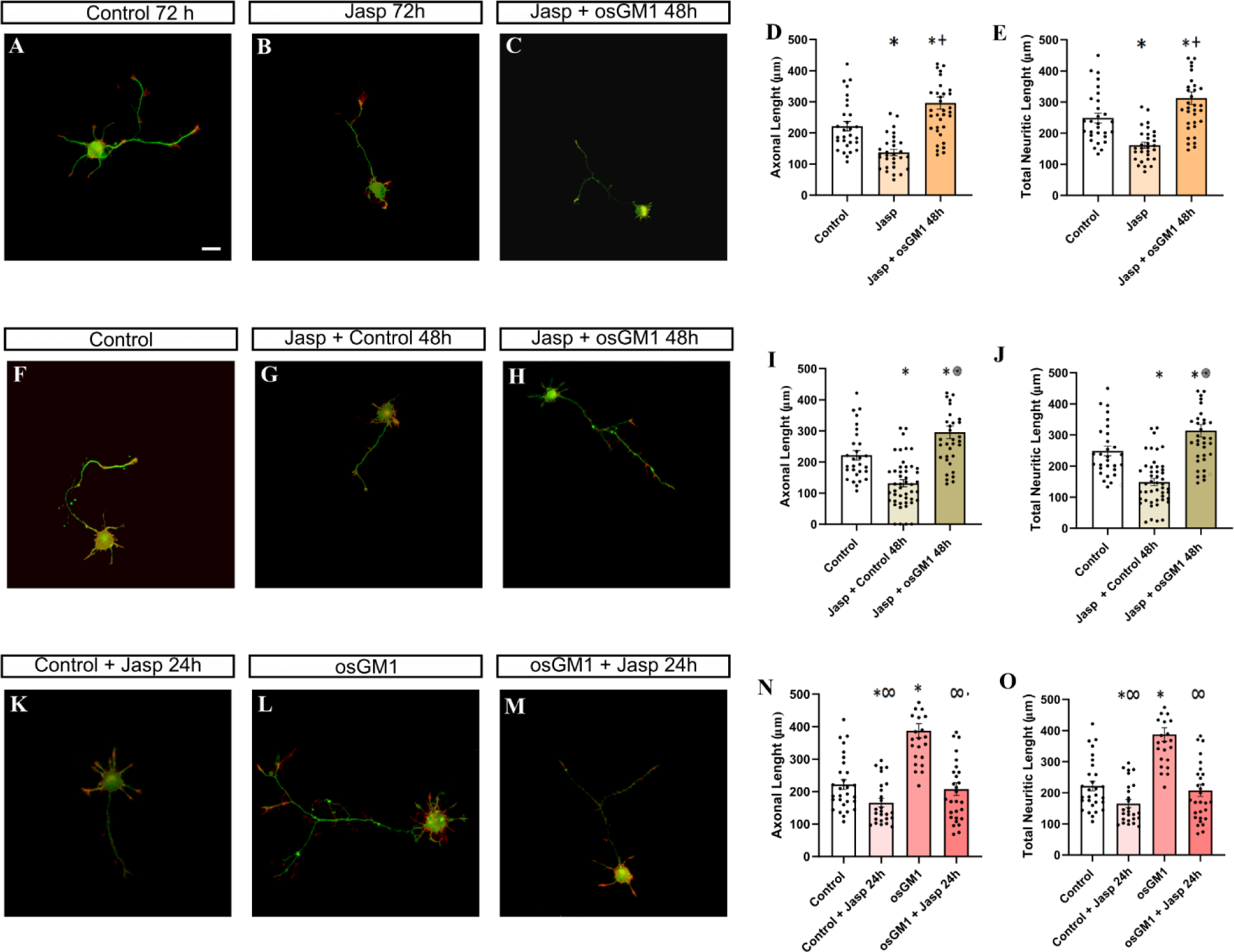
osGM1 reverses actin stabilization induced deficits in axonal elongation and neuritic growth. (A-C, F-H, K-M) Representative images of hippocampal neurons under control conditions or treated with jasplakinolide and/or osGM1 using neuroprotective (A-C, F-H) or neurorestorative paradigms (K-M). Neurons were stained with mAb against tyrosinated α-tubulin (green) and phalloidin-Rhodamine (red). Scale bar: 20 μm. (D, E; I, J; N, O) Graphs showing quantification of o axonal length (D, I, N) and total neurite length (E, J, O) per neuron under the corresponding treatment conditions. Prolonged actin stabilization by jasplakinolide impaired axonal elongation and neuritic growth, whereas osGM1 reversed or attenuated these deficits, consistent with restoration of actin dynamics required for neuronal polarization. For all quantitative analyses, 25–45 neurons were analyzed per treatment condition. Graphs represent mean ± SEM. Statistical analysis: one-way ANOVA followed by Duncan’s post hoc test (p< 0.05). Significant differences versus control (*), jasplakinolide-treated (+), osGM1-treated (∞), and jasplakinolide + control at 48 h (#).

Because these results suggested a restorative action of osGM1 even when actin was already stabilized, we next evaluated whether early ganglioside exposure confers protection against later cytoskeletal alterations. We compared control neurons (Fig. 2F), neurons exposed to Jasplakinolide during the first 24 h followed by 48 h in fresh medium (Fig. 2G), and neurons exposed to Jasplakinolide during the first 24 h and subsequently treated with osGM1 during the last 48 h (Fig. 2H). ANOVA again revealed significant differences in axonal length (F(2,108) = 30.36, p ≤ 0.0001) and total neuritic length (F(2,108) = 31.65, p ≤ 0.0001) (Fig. 2F-J). In this paradigm, the application of jasplakinolide during the first 24 h significantly reduced both axonal length and total neuritic length compared with controls, an effect that persisted even after 48 h in fresh medium, consistent with previous studies describing the long-lasting consequences of excessive actin stabilization (Bubb, Spector, Beyer, & Fosen, 2000; Lazaro-Dieguez et al., 2008; Merriam et al., 2013). In contrast, the presence of osGM1 during the last 48 h reversed these deficits. It restored values to levels even higher than those of controls, indicating a robust restorative effect when the ganglioside acts before cytoskeletal alterations become consolidated.

Finally, to determine whether the action of osGM1 persists even when an additional perturbation is introduced during the final phase of the protocol, we compared control neurons maintained for 72 h (Fig. 2A, F), neurons exposed to jasplakinolide only during the last 24 h (Fig. 2K), neurons treated with osGM1 throughout the 72-h incubation (Fig. 2L), and neurons treated with osGM1 and co-exposed to jasplakinolide during the last 24 h (Fig. 2M). ANOVA revealed significant differences in axonal length (F(3,105) = 27.40; p ≤ 0.0001) and total neuritic length (F(3,105) = 27,017; p ≤ 0.0001) (Fig. 2K-O). In this paradigm, continuous exposure to osGM1 during the 72-h incubation significantly increased both axonal and total neuritic lengths, whereas application of jasplakinolide during the last 24 h of culture significantly reduced both axonal and total neuritic lengths. Moreover, when osGM1 was present from the beginning, and jasplakinolide was introduced only during the last 24 h, the ganglioside substantially mitigated the inhibitory effects of the actin stabilizer, partially restoring both axonal elongation and total neuritic length.

Taking together, these results demonstrate that osGM1 not only counteracts rotenone-induced alterations but also reverses deficits arising from excessive actin stabilization. These findings raise the possibility of a direct modulation of the RhoA/ROCK pathway, a central regulator of actin cytoskeletal dynamics.

### osGM1 reverses rotenone-induced activation of RhoA and ROCK

Considering that the RhoA/ROCK pathway tightly controls actin remodeling and considering the strong restorative effects of osGM1 on neuronal polarization and cytoskeletal dynamics, we examined whether these actions involve a direct modulation of RhoA/ROCK signaling. GM1 binds with high affinity to the TrkA receptor and functionally modulates Rho GTPases (Mutoh, Tokuda, Miyadai, Hamaguchi, & Fujiki, 1995) and, given that TrkA activation engages Rho GTPase-dependent signaling pathways(Kurokawa, Nakamura, Aoki, & Matsuda, 2005), GM1 may indirectly influence Rho GTPase activity downstream of TrkA signaling. TrkA activation promotes PI3K/Akt signaling, which indirectly decreases RhoA activity through Rac1 activation and cytoplasmic translocation, thereby preventing RhoA-ROCK activation, a key event required for neurite outgrowth (Nusser, Gosmanova, Zheng, & Tigyi, 2002; Zhu et al., 2015). Accordingly, Chiricozzi and colleagues (Chiricozzi et al., 2017) demonstrated that osGM1 promotes neurite outgrowth via TrkA-dependent PI3K/Akt activation.

To evaluate this hypothesis, RhoA and ROCK activities were measured using the FRET biosensors RhoA2G and Eevee-ROCK (Bisbal et al., 2018; Fritz et al., 2013; C. Li et al., 2017; Wojnacki et al., 2024).

Neurons were electroporated and, 24 h later, treated with DMSO, rotenone, osGM1, or their combinations, as described in Materials and Methods section and radiometric FRET maps were analyzed. Fluorescence images acquired in the CFP channel confirmed a homogeneous distribution of the biosensor across all experimental groups.

A one-way ANOVA revealed significant differences in RhoA activity among the experimental groups (F(4,117) = 16.55, p ≤ 0.0001). Rotenone significantly increased RhoA activity compared to control conditions (Fig. 3A, B, F), consistent with evidence indicating that the toxin activates the Lfc/RhoA/ROCK pathway, thereby blocking axon formation (Bisbal et al., 2018). In contrast, osGM1 alone did not alter basal activity levels relative to control (Fig. 3A, C, F). Moreover, osGM1 applied either before (Fig. 3D, F) or after rotenone treatment (Fig. 3E, F) significantly reduced this activation.

**Figure 3.**
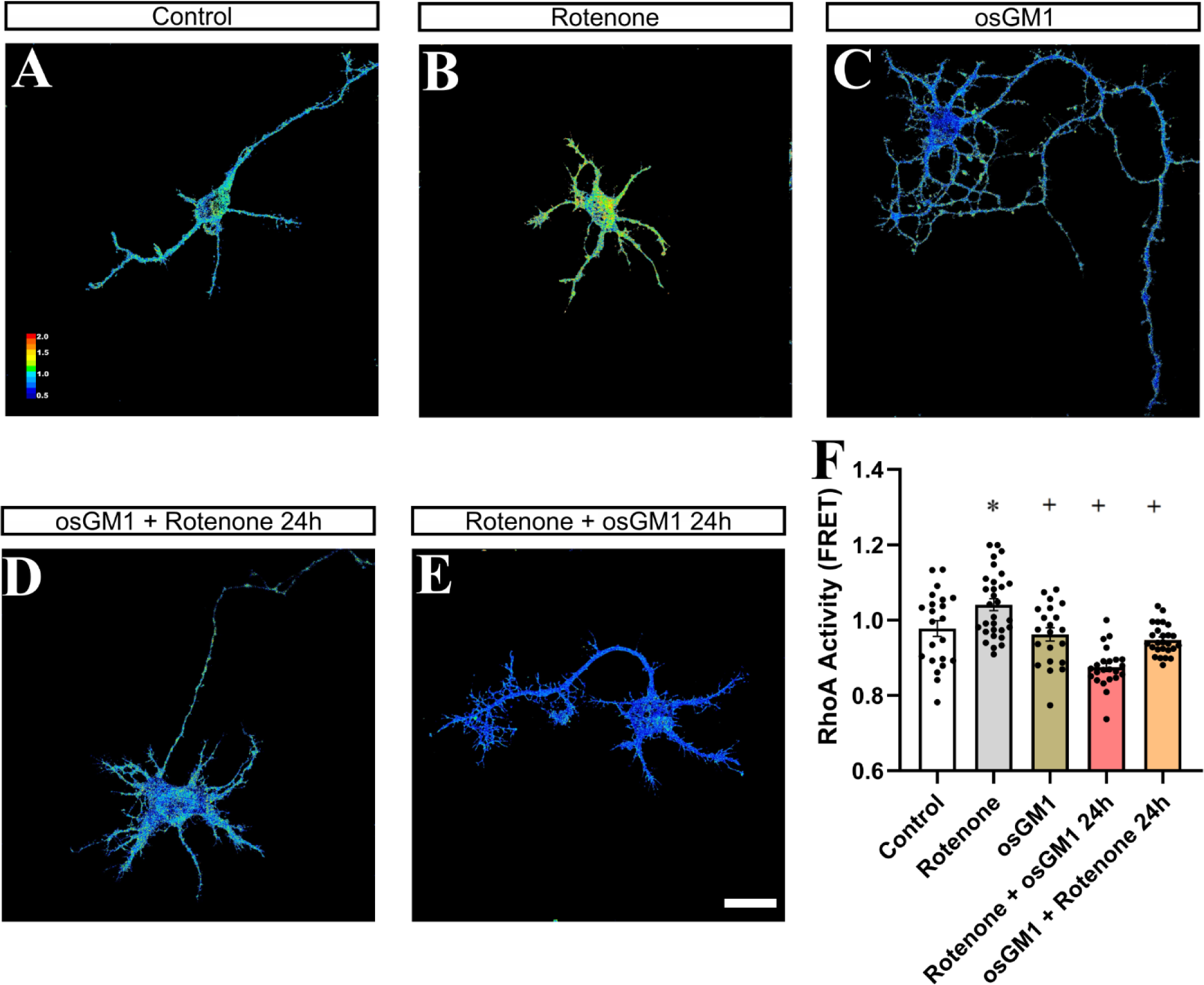
osGM1 reverses rotenone-induced RhoA activation in hippocampal neurons. (A-E) Representative FRET map showing RhoA activity in control neurons and neurons treated with rotenone, osGM1, or their combinations. Scale bar: 20 µm. FRET maps are color-coded according to activation intensity. (F) Graph showing the quantification of RhoA activity in control and treated neurons. Rotenone increased RhoA activity, whereas osGM1, applied either before or after toxin exposure, restored RhoA activity to near-control levels without affecting basal RhoA activity. For all quantitative analyses, 21–30 neurons were analyzed per treatment condition. Graphs represent mean ± SEM. Statistical analysis: one-way ANOVA followed by Duncan’s post hoc test (p < 0.05). Significant differences versus control (*) and rotenone-treated (+).

Given its role as a direct effector of RhoA and a mediator of many of its effects on cytoskeletal dynamics, we next assessed whether alterations in ROCK activation mirrored the changes observed in RhoA activity. A one-way ANOVA showed significant differences in ROCK activity among experimental groups (F(4,107) = 14.75, p ≤ 0.0001). Rotenone significantly increased ROCK activation (Fig. 4B, F) compared with control conditions (Fig. 4A, F), consistent with its ability to activate the RhoA/ROCK pathway. In contrast, osGM1 applied either before (Fig. 4D, F) or after rotenone (Fig. 4E, F) significantly reduced this activation. As observed for RhoA, osGM1 alone did not modify basal ROCK activity levels.

**Figure 4.**
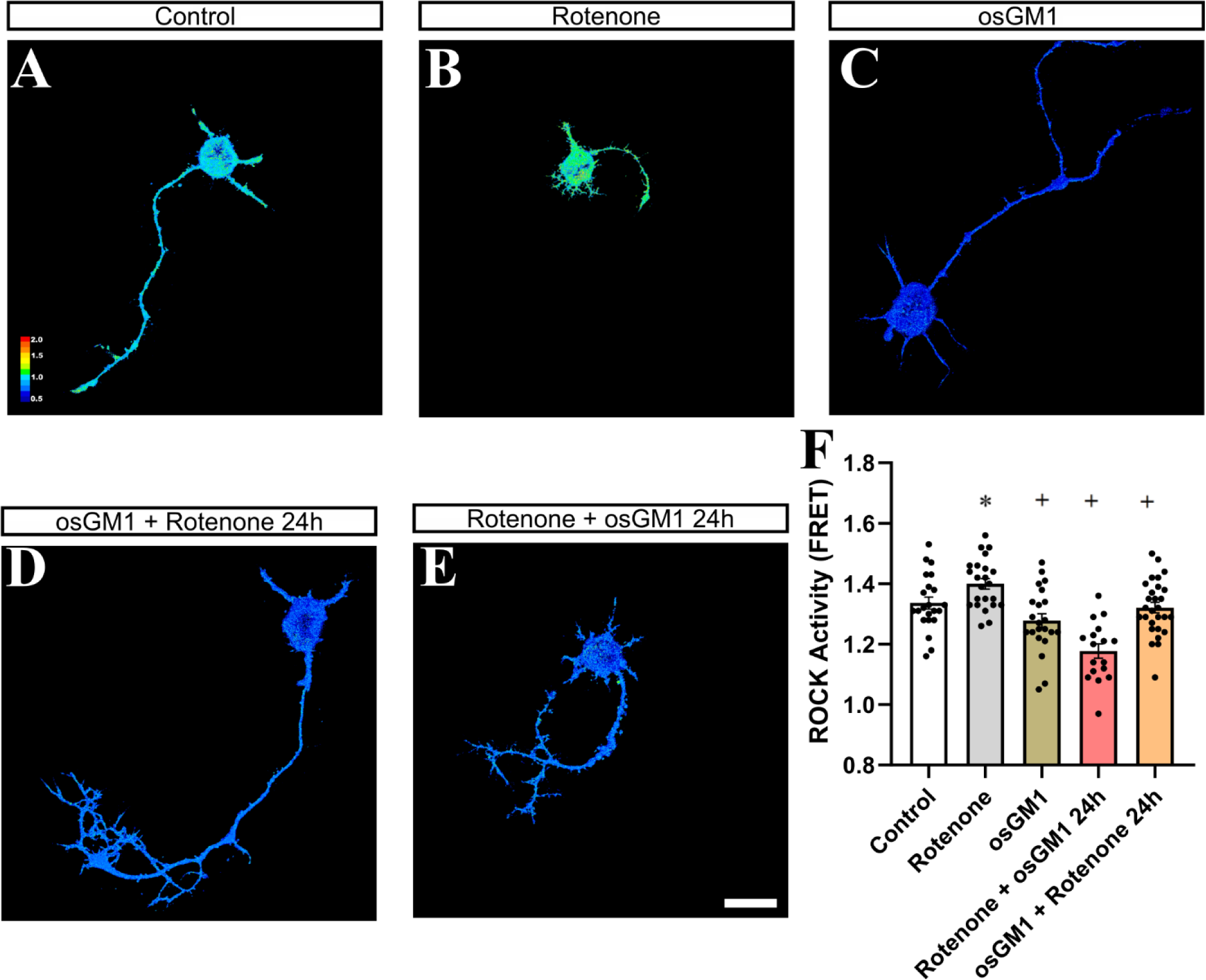
osGM1 attenuates rotenone-induced ROCK activation in hippocampal neurons. (A–E) Representative FRET map showing ROCK activity in control neurons and neurons treated with rotenone, osGM1 or their combinations. Scale bar: 20 µm. FRET maps are color-coded according to activation intensity. (F) Graph showing the quantification of RhoA activity in control and treated neurons. Rotenone induced robust ROCK activation, which was significantly attenuated by osGM1 under both neuroprotective and neurorestorative conditions. For all quantitative analyses, 17–28 neurons were analyzed per treatment condition. Graphs represent mean ± SEM. Statistical analysis: one-way ANOVA followed by Duncan’s post hoc test (p < 0.05). Significant differences versus control (*) and rotenone-treated (+).

Finally, given that actin dynamics -which are tightly regulated by the RhoA/ROCK pathway- are critical for mitochondrial distribution, positioning, and network organization in neurons (Ji, Hatch, Merrill, Strack, & Higgs, 2015; Rehklau et al., 2017), the following section evaluates whether rotenone-induced cytoskeletal disruption extends to the mitochondrial network and whether osGM1 can attenuate these deficits.

### Effects of Rotenone and osGM1 on Mitochondrial Morphology and Function in Hippocampal Neurons

Given that the RhoA/ROCK pathway tightly regulates actin cytoskeleton dynamics, and that actin organization critically influences mitochondrial distribution, positioning, and network maintenance in neurons (Ji et al., 2015; Rehklau et al., 2017), we examined whether rotenone-induced cytoskeletal alterations extend to mitochondrial architecture. Moreover, because mitochondrial morphology is intimately coupled with organelle function -including membrane potential, bioenergetic integration, and cellular stress responses (Eisner, Picard, & Hajnoczky, 2018; Youle & van der Bliek, 2012) we assessed both structural parameters and functional status. In this context and considering osGM1’s ability to protect against maturational arrest and reverse rotenone-induced RhoA/ROCK hyperactivation, we analyzed whether these treatments modify mitochondrial density, morphology, and function in hippocampal neurons.

As shown in Table 1 and Figure 5, the number of mitochondria per cell was significantly reduced in neurons exposed to 0.1 µM rotenone compared with all other groups (F(4,85) = 5.30, p = 0.0007), indicating an adverse effect on mitochondrial density. Similarly, total mitochondrial area was decreased in the rotenone-treated group, whereas neurons treated with osGM1 + rotenone exhibited intermediate values (F(4,85) = 2.20, p = 0.0763). Taken together, these findings indicate that rotenone markedly reduces both mitochondrial number and total mitochondrial area, while osGM1 partially attenuates these effects. This pattern suggests that RhoA/ROCK pathway dysfunction may be accompanied by a progressive dismantling of the mitochondrial network, prompting a more detailed morphological analysis.

**Figure 5.**
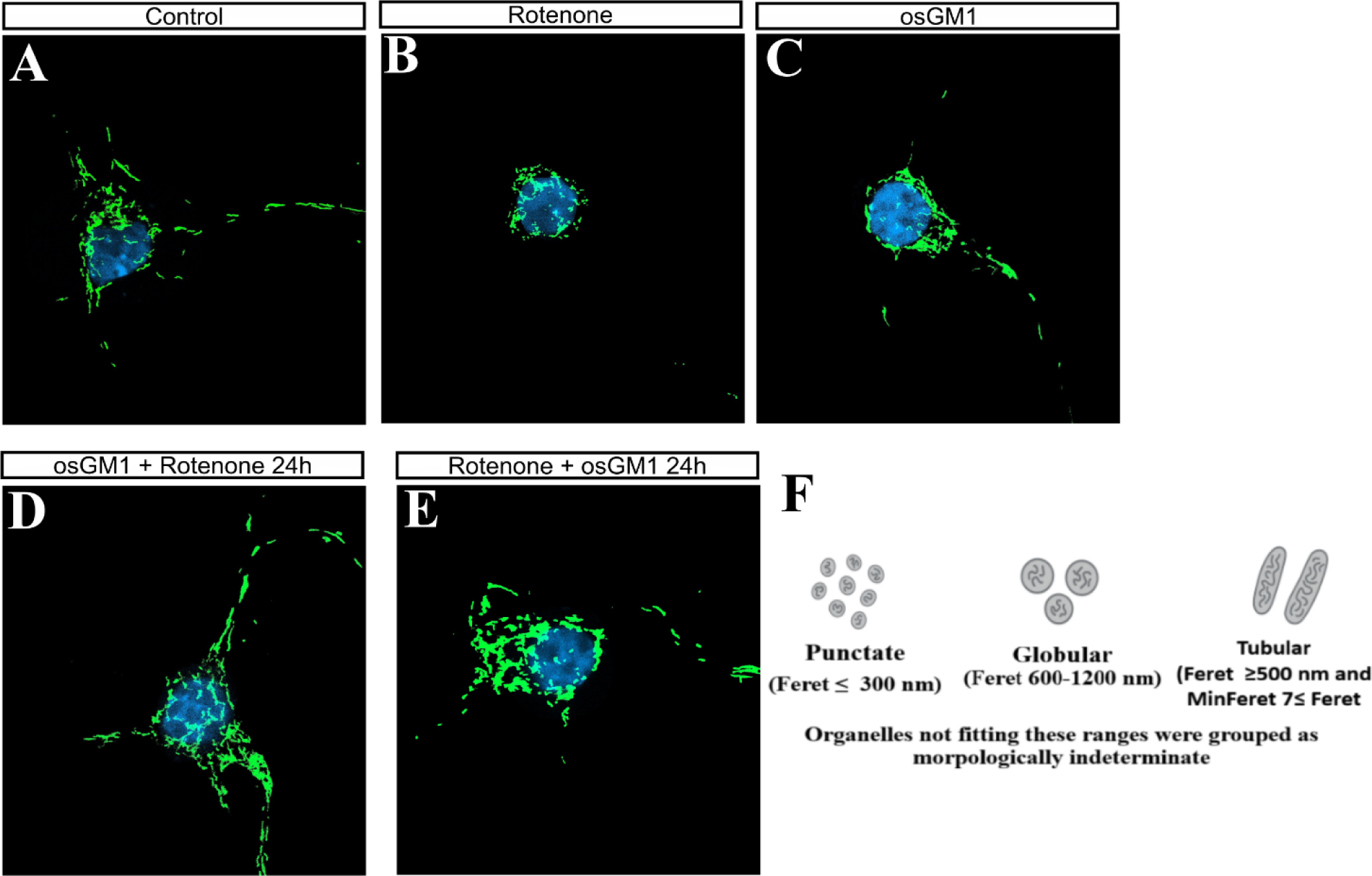
Effects of osGM1 on mitochondrial morphology in hippocampal neurons. (A-E) Representative images of hippocampal neurons expressing MitoYFP (green) and stained with DAPI (blue), to visualize nuclear areas, treated for 48 h with DMSO (Control, A), rotenone (B), osGM1 (C), osGM1 + rotenone (D) or rotenone + osGM1 (E). Images show treatment-dependent alterations in mitochondrial organization and quantified in Table 1. Scale bar: 20 µm. Rotenone markedly reduced mitochondrial density and network complexity, whereas osGM1 alone or in combination with rotenone partially preserved normal mitochondrial morphology. (F) Classification criteria for mitochondrial morphotypes: punctate (Feret ≤ 300 nm), globular (Feret 600–1200 nm), tubular (Feret ≥ 500 nm and MinFeret < 0.7 × Feret), and other irregular or transitional forms (Helguera et al., 2013).

**Table 1.** Effects of treatments on mitochondrial morphometric parameters and morphology in hippocampal neurons. The table shows mean values ± SEM of mitochondrial morphometric parameters, including count, area, perimeter, Feret diameter, and MinFeret in control hippocampal neurons and neurons treated with rotenone, osGM1, or their combinations and Mitocondial morphological categories, mitochondria were classified as punctate (≤ 300 nm), globular (600–1200 nm), tubular (≥ 500 nm and MinFeret < 0.7 × Feret), or indeterminate morphology. The distribution of these categories reflects treatment-dependent changes in mitochondrial size, elongation, and compactness. For all quantitative analyses, 15–16 neurons were analyzed per treatment condition. Statistical analysis was performed using one-way ANOVA followed by Duncan’s post hoc test (p < 0.05) Significant differences versus control (*), rotenone-treated (+), and osGM1-treated (∞) groups are indicated.

| Treatment<br>Parameter | Control | Rotenone | osGM1 | osGM1+Rotenone | Rotenone+osGM1 |
| --- | --- | --- | --- | --- | --- |
|  | **Morphometric Parameters** |  |  |  |  |
| **Count** | 59.40±18.91 | 34.36±11.63* | 74.94±46.35<sup>+</sup> | 81.83±39.94<sup>+</sup> | 62.40±22.87<sup>+</sup> |
| **Area (µm<sup>2</sup>)** | 75.20±25.89 | 52.51±15.12* | 70.19±31.24<sup>+</sup> | 66.40±19.60<sup>+</sup> | 73.93±23.95<sup>+</sup> |
| **Perimeter (µm)** | 6.07±1.45 | 6.55±2.21 | 5.87±2.02 | 5.49±2.40 | 6.52±1.97 |
| **Feret (nm)** | 1422.47±239.78 | 1505.64±445.82 | 1321.67±449.50 | 1370.78±397.04 | 1475.15±259.12 |
| **MinFeret (nm)** | 668.21±209.47 | 545.10±317.67 | 633.62±245.95 | 610.40±242.43 | 626.16±292.51 |
|  | **Mitochondrial morphological categories** |  |  |  |  |
| **% Punctate (≤300 nm)** | 23.42±6.24 | 23.77±9.96 | 23.31±12.60 | 22.78±9.02 | 22.03± 8.49 |
| **% Globular (600–1200 nm)** | 25.27±5.56 | 26.62±8.03 | 26.97±8.59 | 28.46±8.04 | 26.84±6.01 |
| **% Tubular (≥500 nm)** | 31.98±5.66 | 32.81±11.51 | 27.32±10.86 | 22.78±9.02<sup>*</sup> | 33.18±6.66 |
| **% Other** | 19.33±6.47 | 16.81±5.40 | 22.40±6.39<sup>+</sup> | 18.00±3.97<sup>∞</sup> | 17.95±5.68 <sup>∞</sup> |

In contrast to these global alterations, individual morphometric parameters, including perimeter and the Feret and MinFeret diameters (which represent mitochondrial maximal length and thickness and are sensitive indicators of changes in mitochondrial elongation), did not differ significantly among groups (perimeter: F(4,85) = 0.93, p = 0.4517; MinFeret: F(4,85) = 0.43, p = 0.7855; Feret: F(4,85) = 1.57, p = 0.1907) (Table 1), indicating that the average morphology of individual mitochondria remained essentially unchanged despite reductions in mitochondrial number and total area.

Consistent with these observations, Helguera-style morphological classification (Helguera et al., 2013) revealed no significant differences in the proportion of punctate or globular mitochondria. However, a substantial reduction in the frequency of tubular mitochondria was observed (F(4,85) = 5.07, p = 0.0010), indicating a modest but coherent increase in mitochondrial network fragmentation. Neurons treated with osGM1 + rotenone showed the most pronounced decrease in tubular forms, whereas osGM1 alone displayed intermediate values, reflecting a distinct pattern associated with the combined treatment (Table 1).

Interestingly, the “Other” category, which includes irregular or transitional morphologies was significantly increased in osGM1-treated neurons, with intermediate values observed in control cells (F(4,85) = 2.61, p = 0.0414) (Table 1). This pattern may reflect enhanced mitochondrial plasticity promoted by osGM1, potentially associated with active remodeling processes.

Given that mitochondrial morphology is tightly linked to mitochondrial function, we complemented the structural analysis with an assessment of mitochondrial membrane potential using MitoTracker Red CM-H₂XRos. Statistical analysis by one-way ANOVA followed by Duncan’s post hoc test revealed significant differences among experimental groups (F(4,859) = 5.41, p = 0.0003). Neurons exposed to rotenone throughout the entire experimental period exhibited a significant decrease in fluorescence intensity compared with the DMSO-treated control, indicating loss of membrane potential and impaired mitochondrial function (Table 2). In contrast, osGM1 alone did not significantly alter fluorescence intensity, suggesting that it does not compromise basal mitochondrial function. Notably, both pre-treatment and post-treatment with osGM1 during the last 24 h in rotenone-exposed neurons resulted in intermediate mitochondrial fluorescence values between those observed in neurons treated with rotenone alone or osGM1 alone, indicating a partial recovery of mitochondrial membrane potential, consistent with the neuroprotective and neurorestorative actions of osGM1 (Table 2).

**Table 2.** Mitochondrial membrane potential assessed by MitoTracker Red fluorescence in hippocampal neurons under different treatments. Mitochondrial membrane potential assessed by MitoTracker Red fluorescence intensity. The table shows mean values (± SEM) of MitoTracker Red fluorescence intensity expressed as mean gray value in control cells and cells treated with rotenone, osGM1, and their combinations. Changes in fluorescence intensity reflect treatment-dependent alterations in mitochondrial membrane potential. For fluorescence intensity analyses, 150–200 cells were analyzed per treatment condition. Significant differences versus the control group (*) were determined by one-way ANOVA followed by Duncan’s post hoc test (*p < 0.05).

|  | **MitoTracker Red Fluorescence Intensity**<br>**(Mean Gray Value)** |
| --- | --- |
| **Control** | 29.90 ±14.42 |
| **Rotenone** | 25.27±10.46* |
| **osGM1** | 29.42±10.15 |
| **osGM1+ Rotenone** | 27.33±7.47 |
| **Rotenone + osGM1** | 27.23±7.46 |

Together, these results indicate that rotenone not only reduces the total number of mitochondria and their occupied area but also promotes mitochondrial network fragmentation and loss of membrane potential in hippocampal pyramidal neurons. In contrast, osGM1 partially counteracts these effects, preserving both structural and functional parameters of the mitochondrial network, in line with its previously observed neuroprotective and neurorestorative actions on neuronal polarization and axonal growth.

## Discussion

Primary cultures of hippocampal pyramidal neurons have become a widely accepted experimental model for investigating the cellular and molecular mechanisms that regulate neuronal polarization, axonal elongation, and dendritic development(Banker, 2018; Caceres, Ye, & Dotti, 2012). This system allows the controlled recapitulation of the sequential stages of neuronal differentiation, ranging from the initial formation of lamellipodia and undifferentiated neurites to the establishment of a dominant axon and multiple dendrites, reflecting the dynamic reorganization of the cytoskeleton and the acquisition of neuronal polarity. In this context, environmental factors and neurotoxic agents, including pesticides and persistent organic pollutants, have been shown to disrupt these early processes by interfering with both cytoskeletal dynamics and cellular homeostasis, ultimately compromising normal neuronal development (Bisbal & Sanchez, 2019; Ko, Tam, Teixeira, & Frampton, 2020; Luna, Neila, Vena, Borgatello, & Rosso, 2021). This heightened sensitivity during the early stages of differentiation justifies the use of this model to detect subtle alterations in neuronal polarization and growth, making it a particularly suitable tool for dissecting mechanisms of neuronal damage and protection.

On this basis, our results confirm and extend previous observations from our group by demonstrating that exposure to rotenone interferes with progression toward stage 3 of morphological development, significantly reducing total neurite length and altering the balance among distinct stages of neuronal maturation (Bisbal et al., 2018; Bisbal & Sanchez, 2019; Sanchez et al., 2008). Consistent with its established role in neuronal differentiation and survival, GM1 exerted protective and restorative effects on neuronal morphology, acting at both the level of membrane organization and receptor-mediated signaling (Chiricozzi et al., 2017; Mutoh et al., 1995). Notably, osGM1 -the oligosaccharide fraction of GM1- displayed greater efficacy than the native ganglioside in promoting neuronal polarization and axonal elongation. Although receptor-binding parameters or ligand-receptor kinetics were not directly evaluated, its reduced structural complexity may contribute to enhanced functional accessibility to membrane receptors and a more efficient modulation of signaling cascades, thereby potentiating its biological effects (Chiricozzi, 2022; Chiricozzi et al., 2017; Mutoh et al., 1995).

Although TrkA activation was not experimentally manipulated in the present study, substantial prior evidence demonstrates that GM1 and osGM1 functionally interact with this receptor and activate the PI3K/Akt and MAPK pathways, which critically modulate the activity of Rho GTPases (Chiricozzi et al., 2017; Mutoh et al., 1995; Rabin & Mocchetti, 1995). Therefore, TrkA-dependent signaling is proposed here as an interpretative mechanistic framework -rather than a conclusion directly demonstrated in this work- to explain the observed effects of osGM1 on cytoskeletal dynamics and neuronal morphology. Recent in vivo studies have further shown that osGM1 crosses the blood–brain barrier and retains the ability to activate TrkA-mediated signaling, reinforcing its therapeutic relevance (Di Biase et al., 2020; Lunghi et al., 2025).

From a mechanistic perspective, rotenone -a potent inhibitor of mitochondrial complex I- induces sustained activation of the RhoA/ROCK pathway, consistent with its reported effects on cytoskeletal contraction and growth cone collapse under stress conditions (Bisbal et al., 2018; Govek, Newey, & Van Aelst, 2005). Our data show that osGM1 effectively counteracts this aberrant activation, restoring RhoA and ROCK activity to levels comparable to controls and promoting morphological recovery. The use of FRET-based biosensors enabled direct measurement of RhoA and ROCK activity, supporting the interpretation that the observed changes reflect genuine modulation of signaling rather than indirect secondary effects (Bisbal et al., 2018; Fritz et al., 2013; C. Li et al., 2017; Wojnacki et al., 2024). Given the central role of the RhoA/ROCK pathway in regulating actin dynamics, the observed recovery of axonal elongation and neuronal polarization likely reflects the restoration of the cytoskeletal plasticity required for neuronal growth.

In line with this mechanistic framework, both GM1 and osGM1 promoted progression toward stage 3 of neuronal development and significantly enhanced axonal elongation following rotenone exposure. Across all analyzed parameters, osGM1 consistently produced more robust effects than GM1, supporting the notion that the oligosaccharide may represent a key -although not exclusive- component of the biological activity of the GM1 ganglioside. This interpretation is consistent with previous studies demonstrating that osGM1 can induce neuritogenesis through activation of TrkA-dependent PI3K/Akt and MAPK pathways (Chiricozzi et al., 2021; Chiricozzi et al., 2017).

The close correspondence between morphological recovery and cytoskeletal modulation further supports a central role for actin dynamics in the effects of osGM1. In this context, experiments with jasplakinolide confirmed that prolonged actin stabilization is sufficient to negatively affect axonal elongation and neuronal differentiation (Bubb et al., 2000; Lazaro-Dieguez et al., 2008). Rather than relying on static structural analyses of the actin network, the approach adopted here focused on functional manipulation of actin dynamics and on quantifying the resulting morphological consequences, an especially informative strategy during the early stages of neuronal polarization (Bradke & Dotti, 1999; Buck & Zheng, 2002). Notably, osGM1 partially attenuated these deficits even when actin stabilization was already established. Although under certain experimental conditions axonal and neuritic lengths exceeded control values, this increase is interpreted as the release of inhibitory constraints imposed by rotenone-induced aberrant activation of the RhoA/ROCK pathway, rather than as a supraphysiological or aberrant treatment effect.

In parallel with its effects on neuronal architecture, our data show that rotenone induces a marked reduction in mitochondrial density and bioenergetic function in hippocampal pyramidal neurons, as evidenced by decreases in mitochondrial number, total mitochondrial area, and mitochondrial membrane. In contrast, osGM1 treatment attenuated these alterations, partially preserving mitochondrial density and membrane potential, suggesting a functional protective effect without necessarily implying complete restoration of the mitochondrial network.

Although individual mitochondrial morphometric parameters remained essentially unchanged, classification analysis revealed a modest reduction in elongated tubular mitochondria and a relative increase in intermediate or irregular forms. This pattern is consistent with an adaptive remodeling of the mitochondrial network in response to stress, rather than an increase in mitochondrial damage per se, in agreement with current models of mitochondrial plasticity (Eisner et al., 2018; Youle & van der Bliek, 2012). In this context, osGM1 appears to contribute to the stabilization of mitochondrial function and the maintenance of cellular bioenergetics, without providing direct evidence of a marked induction of mitochondrial biogenesis. While molecular regulators of mitochondrial fusion and fission were not directly examined in this study, the combined analysis of density, morphology, and membrane potential provides an integrated functional assessment of mitochondrial status at the cellular level, particularly suited to the objectives of the present work.

Consistent with previous reports indicating that GM1 derivatives improve mitochondrial function and redox balance (Fazzari et al., 2024; Finsterwald, Dias, Magistretti, & Lengacher, 2021; Y. Li et al., 2021), our results provide new evidence that osGM1 simultaneously protects mitochondrial structure and function in hippocampal neurons exposed to rotenone. Similar protective effects have been described in other experimental models subjected to complex I-targeting toxins (Fazzari et al., 2020), reinforcing the notion of a conserved mechanism of action.

Taken together, these findings support a model in which osGM1 exerts neuroprotective and neurorestorative effects through the convergence of complementary mechanisms acting on key processes of early neuronal differentiation. By modulating RhoA/ROCK signaling and actin cytoskeletal dynamics, osGM1 promotes neuronal polarization and axonal elongation, while concurrently preserving mitochondrial function and cellular bioenergetics. Although these results are based on an in vitro model, they provide relevant insight into early cellular mechanisms potentially involved in neuropathological conditions and support the translational interest of osGM1 as a modulator of early pathogenic events, particularly those associated with cytoskeletal and mitochondrial dysfunction (Di Biase et al., 2020).

## Declaration of competing interest

The authors declare no competing financial interest.

## Acknowledgments

This work was supported, in part, by grants FONCYT (PICT 2019-2679), CONICET (PIP 2023-2025), Alzheimer’s Association (AARGD 22-973030), and SeCyT-UNC, Argentina. M.S.S., A.P., G.Q., P.H., M.B., S.B.R., M.E.M., and G.D.F. are members of the Scientific National Research of CONICET, Argentina. P.E.A.R. is a member of the research career from the government of Córdoba, Argentina. E.G.Q. is a fellowship holder from PICT 2020-01225, FONCYT. GNC is a fellowship holder from PICT 2019-2679, FONCYT. The authors thank Laura Montroull for technical assistance with cell cultures; Romina Maiorano, Eliana Martinez, Jesica Piovano, Milagros Nigro, and Marisa Gigena for technical assistance with animals from the vivarium; and Silvina Ferrer for administrative support. The authors give a special thanks to TRB Pharma S.A. (Buenos Aires, Argentina) and TRB Chemedica International (Genève, Switzerland) for the generous gift of GM1 ganglioside and GM1 oligosaccharide and for their interest in financing our research project on the effect of GM1 oligosaccharide on axonal growth and neuronal polarity in hippocampal pyramidal neuron cultures. All microscopes used in this work belong to the “Centro de Microscopía Óptica y Confocal Avanzada de Córdoba” (CEMINCO), integrated into the “Sistema Nacional de Microscopía (SNM-MINCyT), Argentina.

## Declaration on the use of generative AI and AI-assisted technologies in the writing process

During the preparation of this work, the author(s) used ChatGPT (OpenAI) to refine writing, check grammar, suggest alternative phrasings, and look for primary sources. Additionally, the Grammarly program was used to correct grammatical and spelling errors, thereby improving the text’s clarity and coherence. After using this tool, the author(s) reviewed and edited the content as needed and took full responsibility for the publication’s content.

## Notes

### Competing Interest Statement

The authors have declared no competing interest.

